# Astrocyte-derived PTPRZ1 regulates astrocyte morphology and excitatory synaptogenesis

**DOI:** 10.1101/2025.04.30.651514

**Authors:** Alex R. Eaker, Hayli E. Spence-Osorio, Katherine T. Baldwin

**Affiliations:** Neuroscience Center, University of North Carolina, Chapel Hill, USA; Department of Cell Biology and Physiology, University of North Carolina at Chapel Hill, USA

**Keywords:** astrocyte, synapse, morphology, development

## Abstract

Protein tyrosine phosphatase receptor type Z1 (PTPRZ1) is one of the most abundantly expressed and enriched proteins in astrocytes during development, yet its function in astrocytes is unknown. Using an astrocyte-neuron co-culture system, we found that knockdown of *Ptprz1* in astrocytes significantly impaired astrocyte branching morphogenesis. To investigate the function of PTPRZ1 in astrocytes during brain development, we generated a *Ptprz1* conditional knockout mouse and deleted *Ptprz1* from astrocytes postnatally, after the bulk of astrogenesis is complete. At postnatal day 21, we found defects in astrocyte morphology and a reduction in excitatory synapse number across multiple layers of the visual cortex, suggesting important functions for astrocytic PTPRZ1 in both astrocyte morphogenesis and synaptogenesis. PTPRZ1 is expressed in several neural cell types, including radial glial stem cells and oligodendrocyte progenitor cells (OPCs), and regulates critical aspects of neurodevelopment, including neurite outgrowth, neuronal differentiation, myelination, and extracellular matrix (ECM) development. Moreover, altered PTPRZ1 expression is associated with schizophrenia and glioblastoma. Therefore, this mouse model is a valuable resource for investigating cell-type-specific PTPRZ1 function in numerous neurodevelopmental and neuropathological mechanisms.

## Introduction

Astrocytes are morphologically complex glial cells that control many critical aspects of central nervous system function, including synapse formation, neurovascular coupling, and ion and fluid homeostasis. In the mouse cortex, the bulk of astrocyte morphogenesis occurs during the second and third postnatal weeks, coinciding with a period of peak synaptogenesis^1,2^. Astrocytes express numerous membrane-bound proteins in a developmentally regulated manner to interact with both secreted factors and cell adhesion molecules. These interactions facilitate bidirectional communication between astrocytes and neurons and promote structural and functional development in both astrocytes and neuronal synapses^2–4^. While the list of molecules involved in astrocyte development continues to expand, substantial knowledge gaps still exist and the function of many proteins that are highly expressed in astrocytes during development is unknown.

*Ptprz1* is one of the most abundantly expressed genes in cortical astrocytes during development and is strongly enriched in astrocytes compared to all other brain cell types^5,6^; however, its function in astrocytes is unknown. *Ptprz1* encodes for <u>p</u>rotein tyrosine <u>p</u>hosphatase receptor type <u>Z1</u> (PTPRZ1), a member of the receptor-like protein tyrosine phosphatase family. Three splice isoforms of PTPRZ1 have been characterized, including membrane-bound long and short isoforms, and a secreted isoform known as phosphacan^7^. Both the long isoform and phosphacan are chondroitin sulfate proteoglycans (CSPG), whereas the short isoform has been detected in both CSPG and non-CSPG form^7,8^. Immunoblotting of rodent brain lysates indicate that all three isoforms are expressed during development^8–10^, and transcriptomic data and histological experiments confirm developmental expression in radial glial stem cells (RGCs)^11^, neurons^12^, and oligodendrocyte precursor cells (OPCs) ^5,6^, albeit at lower levels than astrocytes^5,6,11^.

Prior studies have revealed important functional roles for PTPRZ1 in multiple aspect of neurodevelopment. PTPRZ1 regulates cortical neuron migration in primary culture^13^ and Purkinje cell dendrite morphology in organotypic slice cultures^14^. A number of studies have found key roles for PTPRZ1 in OPC proliferation and differentiation^15–17^, as well as recovery from demyelinating lesions^18,19^. More recently, studies with constitutive *Ptprz1* knockout mice found that loss of *Ptprz1* impairs angiogenesis^20^, and perineuronal net (PNN) structure^21,22^. Importantly, changes in PTPRZ1 expression are linked to schizophrenia^23–25^, and PTPRZ1 is emerging as a therapeutic target for the treatment of glioblastoma^26–29^, substance use disorder^30,31^, and neurodegenerative conditions such as multiple sclerosis^32^ and Alzheimer’s Disease^33^. Though studies have suggested a role for astrocyte-expressed PTPRZ1 in neuronal function^14,34^ and demyelination^35^, the role of astrocytic PTPRZ1 in the aforementioned neurodevelopmental processes and neurological disorders has not been investigated. Moreover, it is unclear whether and how PTPRZ1 is important for proper development and function of astrocytes themselves.

Parsing out the cell-type-specific contributions of PTPRZ1 is challenging, due to its developmental expression in multiple cell types. To overcome this technical limitation and investigate the role of astrocyte-derived PTPRZ1 in postnatal brain development, we generated a new transgenic mouse line with a floxed *Ptprz1* allele and deleted *Ptprz1* from astrocytes postnatally. Our findings reveal roles for astrocytic PTPRZ1 in astrocyte morphogenesis and synaptogenesis and demonstrate the utility of this mouse model for investigating cell-type-specific functions of PTPRZ1.

## Results

### PTPRZ1 regulates astrocyte morphogenesis *in vitro*

Previously published transcriptomic studies identified *Ptprz1* as one of the most abundantly expressed genes in astrocytes isolated from postnatal day 7 (P7) mouse cortex (**Fig 1A**)^5^. *Ptprz1* is strongly enriched in astrocytes compared to all other brain cell types (**Fig S1A**) and its highest expression levels occur during development, though expression remains high throughout the lifespan^36,37^. These same expression and enrichment patterns are observed in astrocytes acutely isolated from human brain tissue (**Fig S1B**)^6^. To investigate the function of PTPRZ1 in astrocytes, we first used primary rat astrocyte and neuron co-cultures to determine whether PTPRZ1 is required for astrocyte morphogenesis *in vitro*. We used short hairpin RNA (shRNA) under control of an hU6 promoter (hU6-shRNA) to knock down *Ptprz1* (shPtprz1) (**Figure S1C**) in primary rat cortical astrocytes and confirmed successful protein depletion via western blot (**Figure S1D**). A scrambled shRNA sequence was used as a control (shScr). To visualize the morphology of *Ptprz1*-depleted astrocytes, we generated plasmids expressing both hU6-shRNA and CAG-GFP (**Figure S1C**), transfected these plasmids into rat primary astrocytes, then co-cultured astrocytes with primary cortical neurons for 48hrs to induce astrocyte ramification. Compared to astrocytes transfected with shScr, shPtprz1 astrocytes showed significantly reduced branching complexity (**Figure 1C-D**), indicating that PTPRZ1 is required for proper astrocyte morphogenesis *in vitro*.

**Figure 1:**
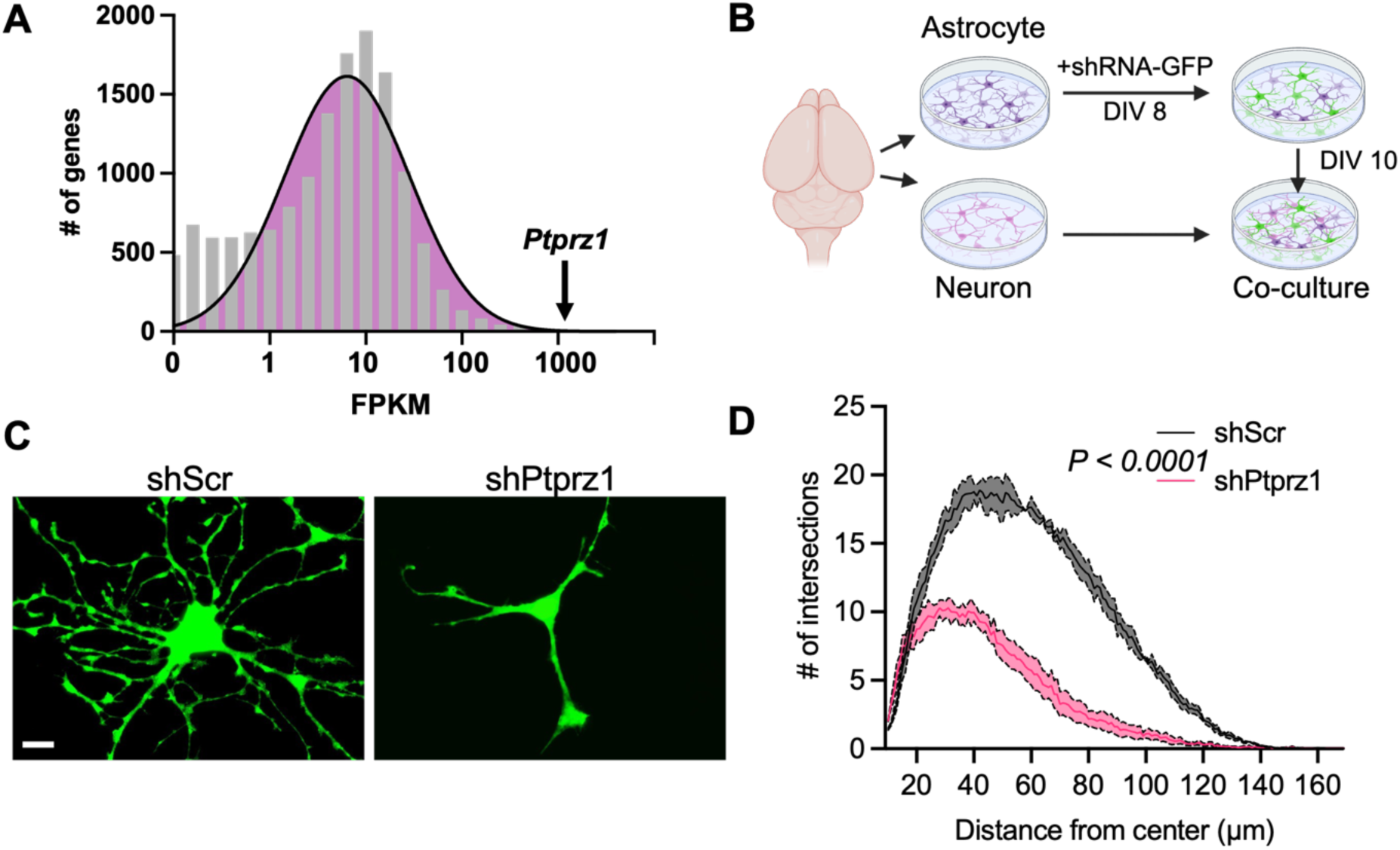
PTPRZ1 regulates astrocyte branching morphogenesis in vitro. **A)** A histogram plot of genes expression levels in purified mouse cortical astrocytes from previously published transcriptomic data^5^. *Ptprz1* ranks tenth. **B)** Workflow for primary astrocyte-neuron co-cultures, created using BioRender.com. **C)** Representative images of astrocytes co-cultured with neurons and expressing scrambled shRNA (shScr) or Ptprz1-targeting shRNA (shPtprz1) and GFP. Scale bar, 10 µm. **D)** Sholl analysis of astrocyte branching complexity. Solid lines represent mean and shaded areas represent SEM, from three independent experiments, >20 cells/condition/experiment, linear mixed model with Tukey HSD.

### Generation of a *Ptprz1* conditional knockout mouse

In our previous studies, we have observed that *in vitro* astrocyte morphology phenotypes often manifest differently *in vivo*^2,38^, likely due to the numerous differences between *in vitro* and *in vivo* microenvironments. Thus, to determine the function of PTPRZ1 in astrocytes during brain development, we generated a transgenic mouse line to conditionally delete *Ptprz1* from astrocytes. To do so, we used a bacterial artificial chromosome with homology to the *Ptprz1* locus to insert *loxP* sites flanking exons 5 and 6 (**Figure 2A**). Cre-mediated excision of Exons 5 and 6 introduces a premature stop codon and is predicted to delete all three PTPRZ1 isoforms.

**Figure 2:**
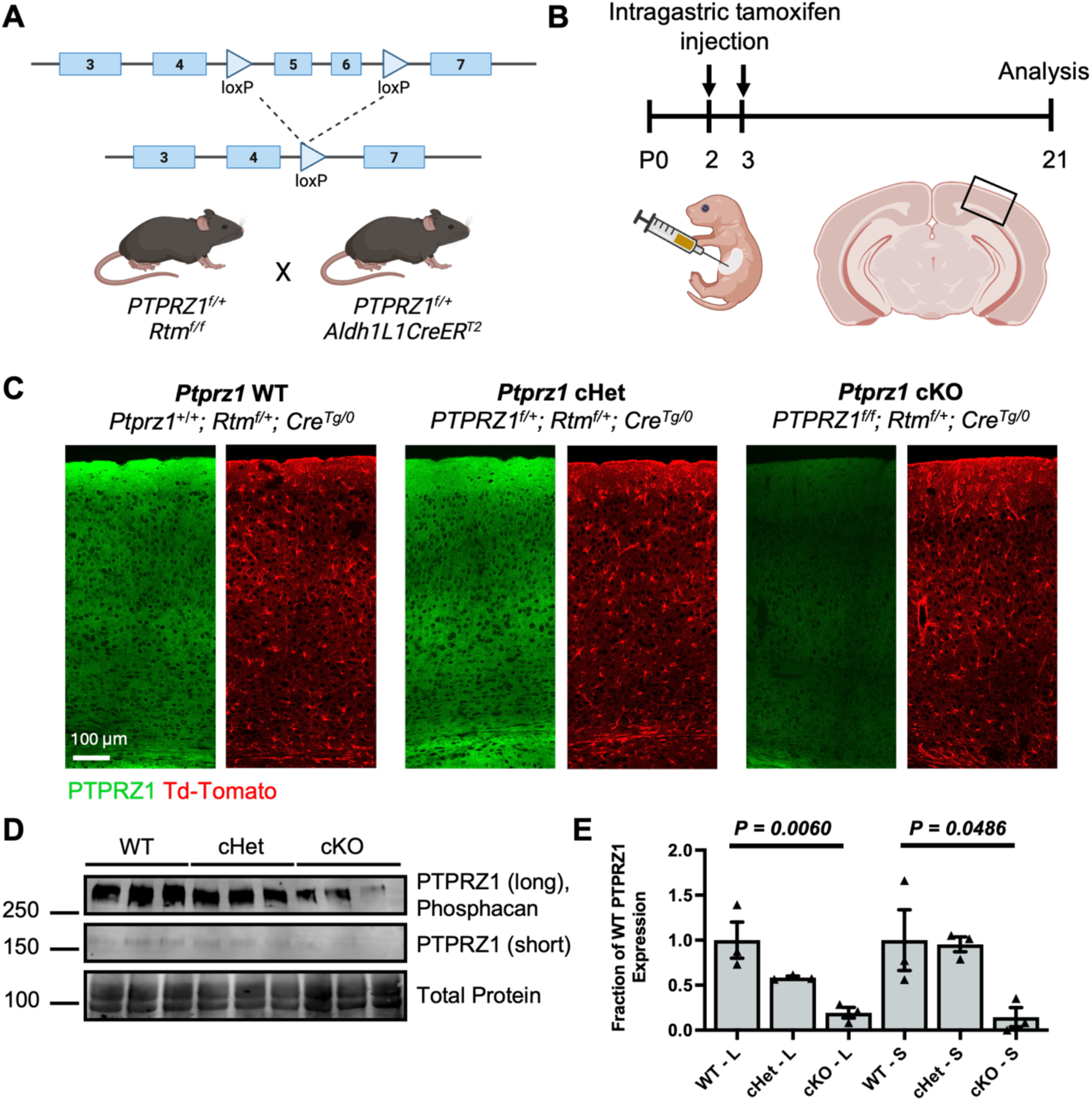
Generation of an astrocyte-specific *Ptprz1* conditional knockout mouse. **A)** Strategy and breeding scheme for conditional deletion of *Ptprz1* from astrocytes, created using BioRender.com. **B)** Experimental timeline for intragastric tamoxifen administration and tissue collection and analysis of the visual cortex, created using BioRender.com. **C)** Representative tile scan images of P21 visual cortex from *Ptprz1* WT, *Ptprz1* cHet, and *Ptprz1* cKO mice with immunolabeling for PTPRZ1 (green) and Td-Tomato (red), demonstrating effective deletion of PTPRZ1 from astrocytes. Scale bar 100 µm. **D)** Western blot image of cortical lysates from 3 WT, 3 cHet, and 3 cKO mice at P21. Lysates were digested with chondroitinase ABC to detect all three isoforms of PTPRZ1. Total protein was used as a loading control. **E)** Quantification of fluorescent band intensity of PTPRZ1 long and phosphacan (L) and PTPRZ1 short (S) normalized to total protein. n = 3 mice per genotype. One-way ANOVA, Dunnett’s post-test.

To delete *Ptprz1* specifically from astrocytes during early postnatal development, we crossed *Ptprz1* flox mice with Aldh1L1CreER^T2^ transgenic mice and administered tamoxifen intragastrically at P2 and P3 (**Figure 2B**). We chose this tamoxifen administration timepoint for three reasons. First, astrocyte-specific deletion with this transgene is unsuccessful prior to astrogenesis, and the majority of astrogenesis in the cortex is complete by P2^39^. Second, PTPRZ1 expression is associated with stemness and pluripotency in human outer RGCs^40^ and suppression of PTPRZ1 promotes OPCs differentiation^15^. Whether PTPRZ1 is involved in astrogenesis is unknown and is complicated by our lack of understanding of the early stages of astrocyte maturation. Deleting *Ptprz1* at the conclusion of astrogenesis eliminates this potentially confounding variable and allows study of astrocyte maturation. Third, we have previously demonstrated that this tamoxifen administration strategy expresses Cre in astrocytes with high specificity and efficiency^38^. To control for any unanticipated phenotypes associated with transgene expression, all mice used in this study expressed exactly one copy of the Cre transgene. These mice also expressed one allele of the Rosa Td-Tomato Cre reporter transgene to visualize Cre-expressing cells.

To confirm successful deletion of PTPRZ1 in astrocytes, we performed immunolabeling of P21 mouse cortex with a Phosphacan antibody (3F8) that detects both the secreted and long isoforms of PTPRZ1. We observed a substantial decrease in PTPRZ1 protein expression in *Ptprz1* conditional knockout (cKO) mice (*Ptprz1^f/f^, RTM^f/+^, Cre^Tg/0^*), but not in *Ptprz1* conditional heterozygous (cHet) mice (*Ptprz1^f/+^, RTM^f/+^, Cre^Tg/0^*), compared to wild type (WT) *Ptprz1^+/+^, RTM^f/+^, Cre^Tg/0^*) (**Figure 2C**). To quantitatively assess the reduction in PTPRZ1 protein levels, we performed western blot of cortical lysates from *Ptprz1* WT, cHet, and cKO mice. Following digest with chondroitinase ABC, all three isoforms were detectable by western blot, though the bands for the long isoform and phosphacan could not be adequately separated for quantification, consistent with prior studies^8^ (**Figure 2D**). We observed an 81% reduction in PTPRZ1-long/phosphacan expression in cKO compared to WT mice, as well as an 86% reduction in PTPRZ1-short expression. PTPRZ1-long/phosphacan expression appeared lower in cHet mice compared to WT, but this difference was not statistically significant (**Figure 2E**). Because we did not observe significant differences in protein expression between WT and cHet mice, we combined these two genotypes for our control group for subsequent experiments, which increased our chances of generating sex-matched littermate pairs for analysis. Collectively, these results demonstrate efficient deletion of *Ptprz1* from astrocytes in the mouse cortex and reveal that the majority of PTPRZ1 protein in the cortex is expressed by astrocytes at this timepoint.

### Postnatal astrocyte-specific deletion of *Ptprz1* does not alter cell number

Astrocytes are known to undergo local division following differentiation in late embryonic and early postnatal stages, with divisions decreasing significantly after the first postnatal week^39^. Because PTPRZ1 is expressed in different stem cell populations and may play roles in cell proliferation and/or differentiation, we wondered whether loss of PTPRZ1 could impact local astrocyte generation following differentiation. To test this idea, we quantified the number of astrocytes in the P21 mouse visual cortex of sex-matched litter pairs of *Ptprz1* control and cKO mice using a new semi-automated workflow (**Figure S3**) based on our previously published nuclear labeling strategy^38^ (**Figure 3A**). At P21, we found no difference in the number of astrocytes (Sox9+/Olig2-) between *Ptprz1* cKO mice and sex-matched littermate controls (**Figure 3B-C**). Neuron (NeuN+) and oligodendrocyte-lineage cell (Olig2+) numbers were also unaffected (**Figure 3D-G**). These results demonstrate that deletion of PTPRZ1 from astrocytes does not impact the later stages of astrogenesis in the visual cortex, nor does it impact the total number of neurons or other Olig2-expressing cells.

**Figure 3:**
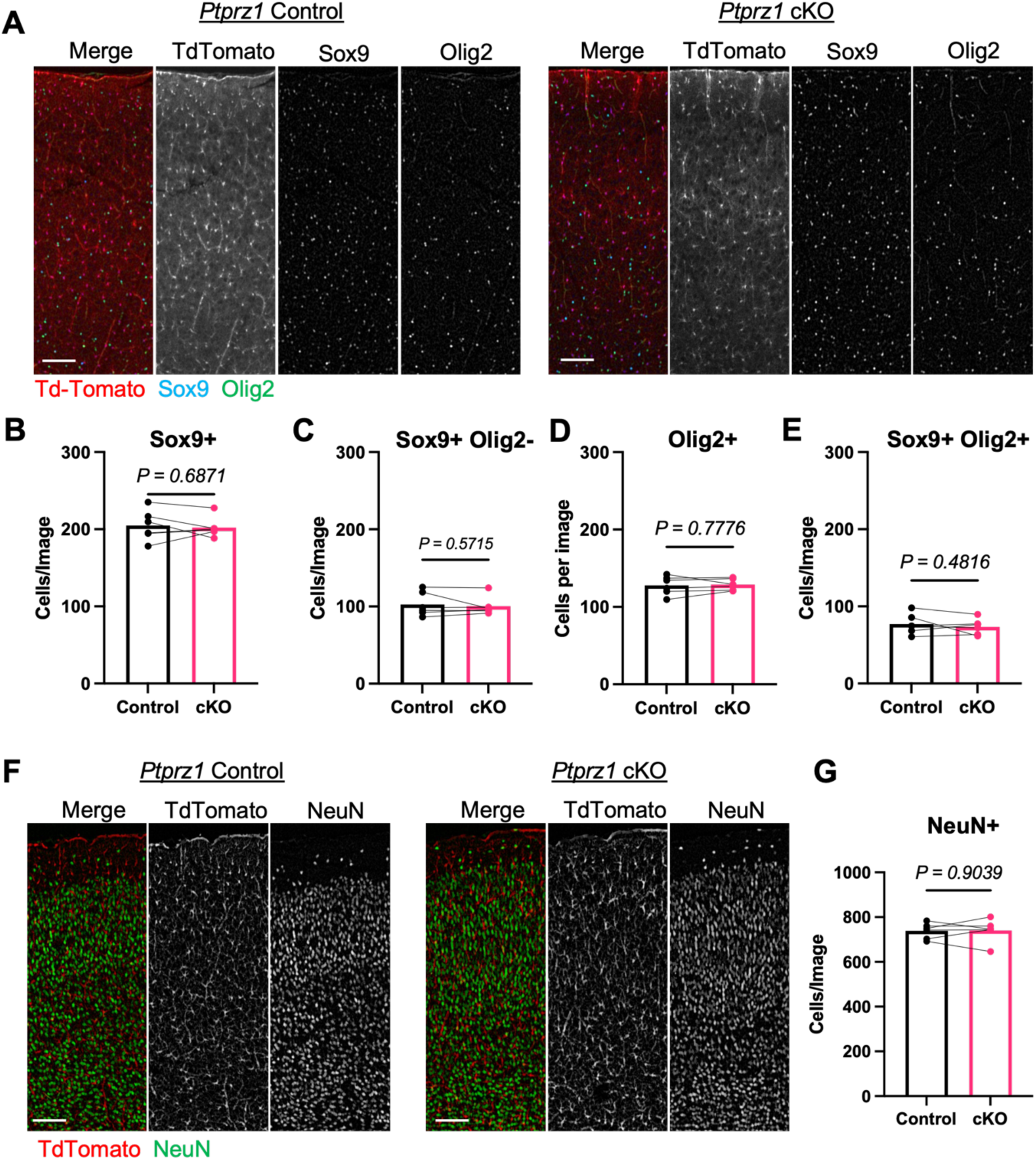
Postnatal deletion of *Ptprz1* from astrocytes does not impact cortical cell numbers. **A)** Representative merged and single-channel tile scan images of P21 visual cortex from *Ptprz1* control (WT and cHet) and *Ptprz1* cKO mice with tdTomato+ (red) astrocytes and immunolabeling for Sox9 (blue) and Olig2 (green). Scale bar 100 µm. **B-E)** Quantification of the number of cells from panel A images for **B)** Sox9+, C) Sox9+/Olig2-(astrocytes), **D)** Olig2+ (oligodendrocyte-lineage cells), and **E)** Sox9+/Olig2+ nuclei. n = 6 sex-matched littermate pairs of control and cKO mice. Lines connect sex-matched control-cKO littermates. Dots represent per animal averages of three images. Paired two-tailed student’s t-test. **F)** Representative merged and single-channel tile scan images of P21 visual cortex from *Ptprz1* control (WT and cHet) and *Ptprz1* cKO mice with tdTomato+ (red) astrocytes and immunolabeling for NeuN (green). Scale bar 100 µm. **G)** Quantification of cells from panel F images with NeuN+ (neurons) nuclei from n = 6 sex-matched littermate pairs of control and cKO mice. Lines connect sex-matched control-cKO littermates. Dots represent per animal averages of three images. Paired two-tailed student’s t-test.

### PTPRZ1 regulates astrocyte morphology *in vivo*

To determine whether *Ptprz1* is necessary for astrocyte morphogenesis *in vivo*, we performed a comprehensive assessment of astrocyte morphology in P21 visual cortex of *Ptprz1* control and cKO mice. To accurately capture the morphology of individual cells, we performed unilateral intracortical injection of PHP.eB serotype adeno-associated virus (AAV) at P1-P2 to sparsely label astrocytes with membrane-targeted GFP (GFP-CAAX) under control of the human minimal GFAP promoter (gfaABC1D) (**Figure 4A**). We acquired confocal z-stack images of entire astrocyte volumes from individual Layer 5 astrocytes and analyzed three-dimensional (3D) astrocyte architecture using our established workflows^41^ (**Figure 4B**). We performed statistical analysis both on a per-cell and per-animal basis. While per-animal analysis conveys the reproducibility amongst independent subjects, per-cell analysis reveals the variability of astrocyte size and morphology within each subject and is the current field-standard^3,42–44^.

**Figure 4:**
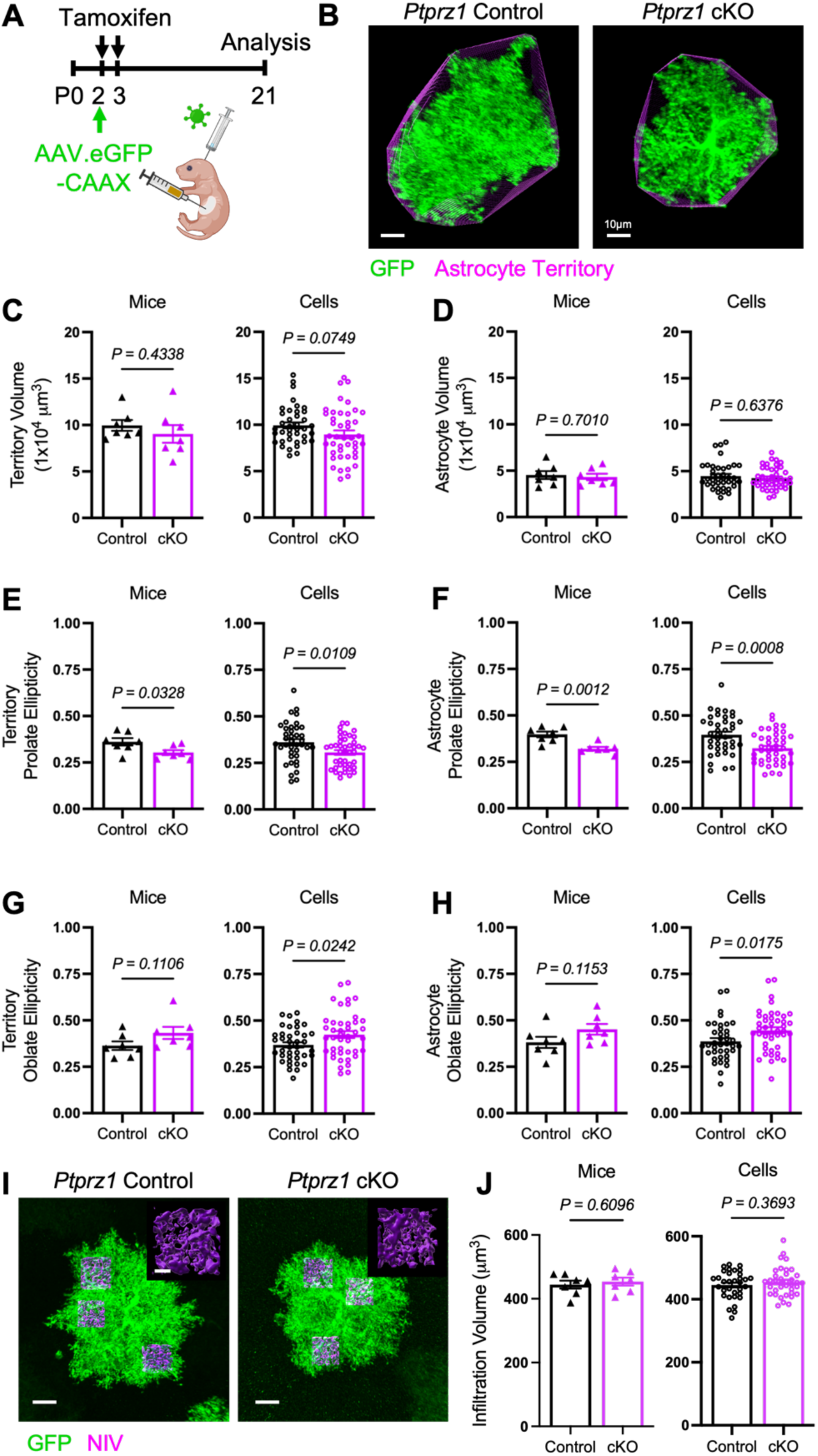
*Ptprz1* cKO astrocytes display altered morphology at P21. **A)** Tamoxifen administration and AAV injection strategy for *Ptprz1* cKO and sparse labeling for detailed morphology analyses. **B)** V1 L5 astrocytes at P21 expressing eGFP-CAAX in *Ptprz1* control (left) and *Ptprz1* cKO (right) mice. Astrocytes expressing eGFP-CAAX in green; astrocyte territory outlined in magenta. Scale bar, 10 µm. **(C-H)** 3D morphology analyses of: **(C)** territory volume, **(D)** astrocyte volume, **(E)** territory prolate ellipticity, **(F)** astrocyte prolate ellipticity, **(G)** territory oblate ellipticity, and **(H)** astrocyte oblate ellipticity in P21 V1 L5 astrocytes. Data presented as animal averages (left; individual mice represented as triangles; n=7 mice per group, 4-8 cells per mouse) and individual astrocyte statistics (right; individual astrocytes represented by open circles). Bars are mean +/- SEM. Unpaired t-test. **I)** Representative V1 L5 astrocytes at P21 expressing eGFP-CAAX (green) with Neuropil Infiltration Volume (NIV) reconstructions (purple; inset scale = 1µm). Scale bar, 10 µm. **J)** NIV analysis for *Ptprz1* control and cKO astrocytes. Three ROIs/cell, 4-5 cells/mouse, 7 mice/condition. Data presented as animal averages and individual astrocyte statistics as above **(C-H).** Bars mean +/- SEM. Unpaired t-test.

Though we observed a dramatic reduction of morphological complexity upon *Ptprz1* knockdown *in vitro*, we did not observe significant differences in astrocyte territory volume (**Figure 4C**), astrocyte cell volume (**Figure 4D**), surface area, or sphericity (**Figure S4A – S4D**) between *Ptprz1* control and cKO astrocyte at P21. We did, however, observe differences in ellipticity of both astrocyte territories and astrocyte cell volumes, with *Ptprz1* cKO astrocytes having a lower prolate ellipticity index (**Figure 4E-F**) and a higher oblate ellipticity index (**Figure 4G-H)** compared to controls. This finding indicates that *Ptprz1* cKO astrocytes are less elongated (prolate) and more flattened (oblate) relative to control astrocytes. Note that ellipticity is measured independently of anatomical orientation and is based on the intrinsic shape of the cell (**Figure S4E**).

To determine whether PTPRZ1 is required for astrocytes to form finer branches that infiltrate the neuropil, we acquired high resolution images from thinner (40 µm) sections and analyzed 3D regions of interest within labeled astrocyte territories to quantify neuropil infiltration volume (NIV) (**Figure 4I**). In contrast to our *in vitro* findings, loss of *Ptprz1* did not impact the formation of finer astrocytes branches, at least at the level of confocal resolution (**Figure 4J**). We also performed an additional multipoint 2-demensional analysis on maximum projections of these same image files, using methods developed in a previous study^42^ and did not identify any significant differences in cell size or complexity (**Figure S4F – S4K**). Collectively these results demonstrate a role for astrocytic PTPRZ1 in regulating astrocyte geometry, with no apparent defects in cell size or complexity visible at confocal resolution at the P21 time point.

### PTPRZ1 regulates excitatory synapse number

Astrocyte development is inextricably linked to synapse development. In the developing mouse cortex, astrocytes promote synapse formation and maturation via secreted factors and direct contact. Multiple studies have detected PTPRZ1 isoforms at the synapse^45,46^, and behavioral studies reveal impaired spatial learning^47^ and contextual fear memory^48^ in constitutive *Ptprz1* KO mice. However, the cell-type-specific roles of PTPRZ1 at the synapse are unknown, and whether any of the PTPRZ1 isoforms are important for synapse development has not been investigated.

To determine whether astrocytic PTPRZ1 is required for proper synapse formation, we quantified the number of excitatory and inhibitory synapses in different layers of the visual cortex from sex-matched littermate pairs of *Ptprz1* cKO and control mice. Because astrocytes control synapse formation with circuit-level precision, we used different excitatory presynaptic markers in combination with excitatory post-synaptic marker PSD95 to differentiate between intracortical (VGlut1/PSD95) and thalamocortical (VGlut2/PSD95) synapses. We used Synbot^49^ to quantify the number of synaptic puncta, defined as the co-localization of pre- and post-synaptic markers. We focused our analysis on layer 1 and layer 4 of the visual cortex, where both excitatory input types are present. We also examined excitatory intracortical synapses in layer 5, where the bulk of our morphology analysis was performed. *Ptprz1* cKO mice showed significant reductions in excitatory intracortical synapse number in layer 1 and layer 5 but not layer 4 (**Figure 5A-F**). Thalamocortical synapse number was reduced in layer 1 and layer 4 in *Ptprz1* cKO mice compared to controls (**Figure 5G-J**). We also examined inhibitory synapse number in layers 1, 4, and 5 by quantifying the co-localization of inhibitory presynaptic marker VGAT and inhibitory postsynaptic gephyrin. In contrast to excitatory synapses, we did not observe any significant difference in inhibitory synapse number across layers (**Figure S5A-F**). Collectively, these results indicate that astrocytic PTPRZ1 is required for normal excitatory synapse development in the mouse visual cortex.

**Figure 5:**
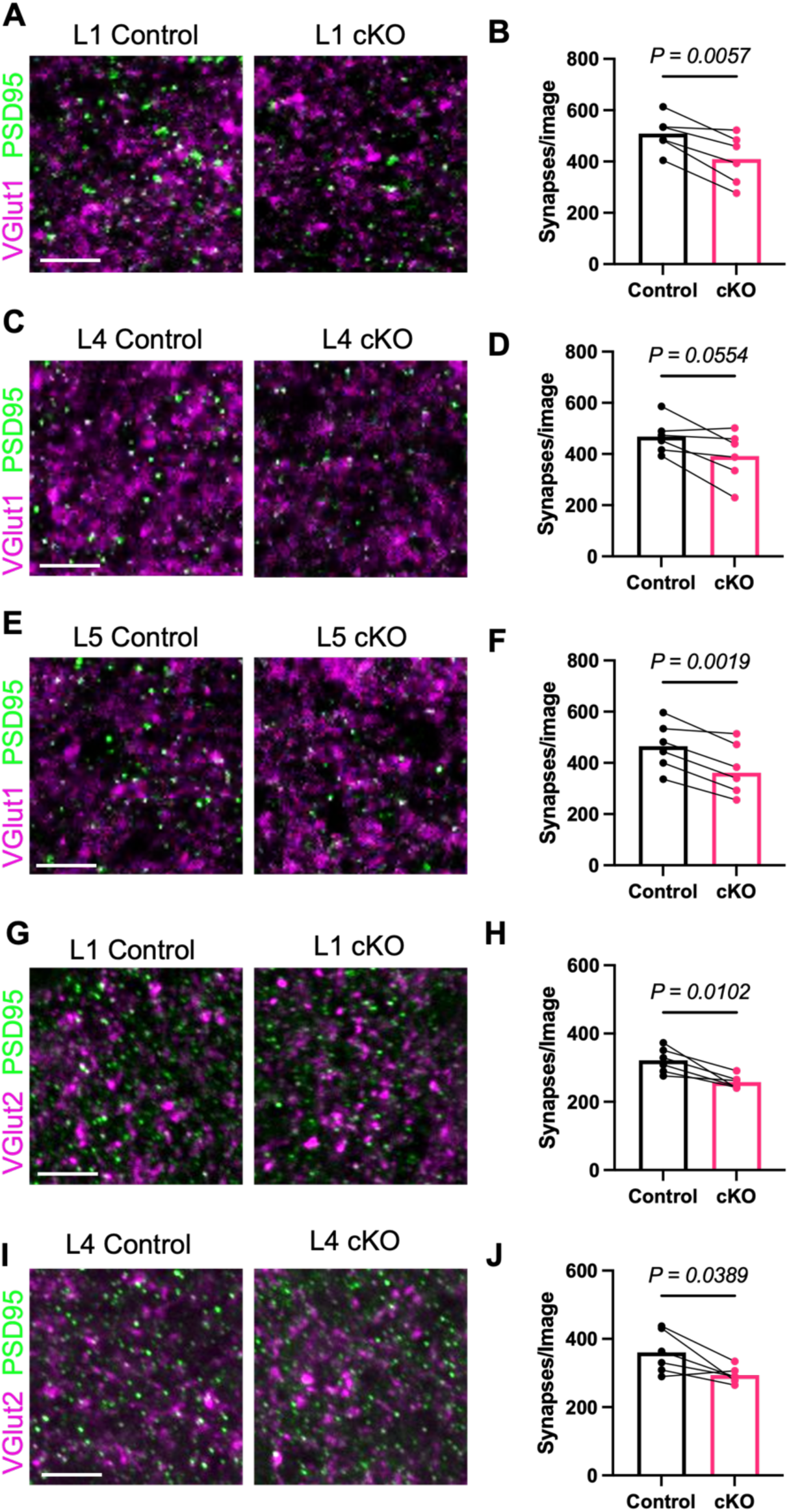
Astrocytic-PTPRZ1 regulates excitatory synapse number *in vivo*. **A)** Representative 20 µm x 20 µm regions of interest (ROIs) labeled with excitatory intracortical synapse markers in L1 visual cortex at P21. Presynaptic VGlut1 (magenta) and postsynaptic marker PSD95 (green). Scale bar 5 µm. **B)** Quantification of number of co-localized VGlut1/PSD95 puncta per image (129.35 µm x 129.35 µm) from L1 visual cortex. **C)** Representative ROIs labeled with VGlut1 and PSD95 in L4 and **D)** quantification of co-localized puncta. **E)** Representative ROIs labeled with VGlut1 and PSD95 in L5 and **F)** quantification of co-localized puncta. **G)** Representative ROIs labeled with excitatory thalamocortical synapse markers in L1 visual cortex at P21. Presynaptic VGlut2 (magenta) and postsynaptic marker PSD95 (green). Scale bar 5 µm. **H)** Quantification of number of co-localized VGlut2/PSD95 puncta per image (129.35 µm x 129.35 µm) from L1. **I)** Representative ROIs labeled with VGlut2 and PSD95 in L4 and **J)** quantification of co-localized puncta. For B, D, F, H, and J: n = 6 sex-matched littermate pairs of control and cKO mice. Lines connect sex-matched control-cKO littermates. Dots represent per animal averages of 15 images. Paired two-tailed student’s t-test.

## Discussion

Here, we developed a new *Ptprz1* conditional knockout mouse to study the astrocyte-specific functions of PTPRZ1 during brain development. We demonstrate successful astrocyte-specific postnatal deletion of all three PTPRZ1 isoforms and uncover roles for astrocytic PTPRZ1 in astrocyte morphogenesis and excitatory synaptogenesis.

In our initial experiments with primary cultures, we observed a dramatic reduction in astrocyte branching complexity upon *Ptprz1* knockdown. However, in P21 *Ptprz1* cKO mice, we observed changes in astrocyte 3D architecture, with *Ptprz1* cKO astrocytes having altered ellipticity compared to control astrocytes, but no changes in astrocyte size or complexity. Several factors could explain this discrepancy between *in vitro* and *in vivo* findings: 1) Given the responsiveness of astrocytes to their microenvironment^42^, genetic manipulations to primary astrocytes in 2-D culture with neurons may produce different phenotypes than genetic manipulations to astrocytes in the 3-D environment of the cortex, as other studies have shown^38^. 2) Sparse *in vitro* knockdown of *Ptprz1* allows for imaging of individually labeled astrocytes. In this scenario, knockdown astrocytes are surrounded by wild-type astrocytes that express PTPRZ1 and may compete with knockdown astrocytes for resources or space. In contrast, our *in vivo* deletion removes PTPRZ1 from all astrocytes. 3) The cultured astrocytes are only in contact with neurons for 48 hours prior to imaging and are younger (DIV 12) than P21 astrocytes *in vivo*. Given that PTPRZ1 expression in the mouse cortex is highest at P7^36^, earlier ages may show a more dramatic morphological phenotype that largely resolves by P21. Future studies are needed to determine whether loss of PTPRZ1 impacts the earlier stages of astrocyte morphogenesis. 4) Lastly, finer astrocyte leaflets are not resolvable by confocal microscopy^1^. If PTPRZ1 regulates development of astrocyte leaflets and perisynaptic processes, super resolution microscopy and/or electron microscopy would be required to visualize this phenotype.

Whether the changes we observed in astrocyte ellipticity are relevant to astrocyte function is unclear. Few studies have examined astrocyte ellipticity as a morphological metric, though a recent study noted changes in astrocyte ellipticity that coincided with transcriptional changes following traumatic brain injury^50^. Visually, the decrease in prolate ellipticity and corresponding increase in oblate ellipticity appears to be driven by the decreased propensity of *Ptprz1* cKO astrocytes to extend their branches outside of spherical arbor. This could be reflective of abnormal engagement of *Ptprz1* cKO astrocytes with their tissue microenvironment. Future studies are necessary to determine whether this is the case and whether alterations to astrocyte ellipticity reflect changes in molecular and/or functional states.

Elucidating the role of astrocytic PTPRZ1 in excitatory synapse formation also requires further investigation. PTPRZ1 may act directly at neuronal synapses to promote synapse formation as a secreted factor (phosphacan) and/or as a transmembrane receptor. Alternatively, lack of PTPRZ1 expression in developing astrocytes may alter astrocyte morphology and/or physiology more broadly, thereby hindering the ability of astrocytes to promote synaptogenesis or stabilize newly formed synapses. Several PTPRZ1 binding partners have been described, including pleiotrophin (PTN), a secreted growth factor that regulates various aspects of nervous system development^51^. Whether any astrocyte-specific PTPRZ1 functions during brain development are PTN-dependent is an exciting topic for future investigation, particularly given the interest in PTN as a therapeutic target to treat nervous system injury^52,53^, alcohol use disorder^54^, and other neurological disorders^30^.

In addition to astrocytes, PTPRZ1 is expressed in various cell types during early nervous system development, including radial glial stem cells and oligodendrocyte precursor cells^5,6,11^. Several studies suggest important functions for PTPRZ1 in numerous aspects of nervous system development and plasticity across multiple cell types, yet the cell-type-specific functions of PTPRZ1 are unknown. This mouse model will be a useful tool for cell-type-specific and temporal deletion of *Ptprz1* to understand its function in healthy brain development. Importantly, PTPRZ1 is emerging as a therapeutic target for a growing list of neurological and neuropsychiatric disorders, including schizophrenia, glioblastoma, and substance use disorder^29–31,55^. Thus, understanding the cell-type-specific functions of PTPRZ1 is essential to develop effective therapeutic strategies to target nervous system dysfunction while mitigating off-target consequences of manipulating PTPRZ1 function.

## Materials and Methods

### Animals

All mice were used in accordance with the Institutional Animal Care and Use Committee (IACUC) and the UNC Department of Comparative Medicine (IACUC Protocol Numbers 21-116.0 and 24-005.0). Mice were housed in standard conditions with 12-hour day/night cycles. Aldh1l1-Cre/ERT2 BAC transgenic (JAX 029655), ROSA-td-Tomato Ai14 (RTM) (JAX 007914), FLPo (JAX 012930), and C57BL/6J (JAX 000664) lines were obtained through Jackson Laboratory.

*Ptprz1* conditional knockout mice were generated using homologous recombination collaboration with the Duke Transgenic Mouse Facility. Briefly, a bacterial artificial chromosome (BAC) with homology to the *Ptprz1* genomic locus was generated to insert *loxP* sites before Exon 5 and after Exon 6. BACs were injected into mouse ES cells (G4-129S6B6F1) and positive clones were injected into pseudo-pregnant females. Chimeric offspring males were mated to C57BL/6J females to achieve germline transmission, identified via PCR amplification of genomic DNA using the following primers: Forward 5’ TCACAAGGGTTAGCTTCACAG 3’ and Reverse 5’ AGCAGTAGACTTGCATCTGTG 3’. (WT = 644 bp, Flox = 733 bp). Mice with germline transmission were mated with *Flpo* transgenic mice on a C57BL/6J background to excise the Neomycin selection cassette. Recombined offspring were mated with C57BL/6J, RTM, or Aldh1l1-CreERT2 transgenic mice to generate breeding pairs for experiments.

Littermate pairs of the same sex were assigned to experimental groups based on genotype and collected for experiments at postnatal day 21 (P21). For all experiments, mice of both sexes were included in analysis. We did not observe any influence or association of sex or litter on the experimental outcomes. Criteria for inclusion, exclusion, and randomization are listed for each experiment in specific method subsections.

### Cell Culture

#### Cortical neuron

Purified rat cortical neurons were prepared as described previously^38^. Briefly, cortices were micro-dissected from P1 rat pups of both sexes, digested in papain (7.5 units/mL), triturated in low and high ovomucoid solutions, resuspended in panning buffer (DPBS (GIBCO 14287) supplemented with BSA and insulin) and passed through a 20 µm mesh filter (Elko Filtering 03-20/14). Filtered cells were incubated on negative panning dishes coated with Bandeiraea Simplicifolia Lectin 1, followed by goat anti-mouse IgG+IgM (H+L) (Jackson ImmunoResearch 115-005-044), and goat anti-rat IgG+IgM (H+L) (Jackson ImmunoResearch 112-005-044) antibodies, then incubated on positive panning dishes coated with mouse anti-L1 (ASCS4, Developmental Studies Hybridoma Bank, Univ. Iowa) to bind cortical neurons. Adherent cells were dislodged using a P1000 pipet, pelleted (11 min at 200 g), and resuspended in serum-free neuron growth media (NGM; Neurobasal, B27 supplement, 2mML-Glutamine, 100U/mL Pen/Strep, 1mM sodium pyruvate, 4.2 µg/mL Forskolin, 50 ng/mL BDNF, and 10 ng/mL CNTF). 70,000 neurons were plated onto 12 mm glass coverslips coated with 10 µg/mL poly-D-lysine (PDL, Sigma P6407) and 2 µg/mL laminin and incubated at 37°C in 10% CO_2_. On day *in-vitro* (DIV) 2, half of the media was replaced with NGM and AraC (10 µM) was added to stop the growth of proliferating contaminating cells. On DIV 3, the media was replaced with NGM. Neurons were fed on DIV 6 and DIV 9 by replacing half of the media with NGM.

#### Cortical astrocytes

Rat cortical astrocytes were prepared as described previously^38^. P1 rat cortices from both sexes were micro-dissected, digested in papain, triturated in low and high ovomucoid solutions, filtered, and resuspended in astrocyte growth media (AGM; DMEM (GIBCO 11960), 10% FBS, 10 µM hydrocortisone, 100 U/mL Pen/Strep, 2 mM L-Glutamine, 5 µg/mL Insulin, 1 mM Na Pyruvate, 5 µg/mL N-Acetyl-L-cysteine). Between 15-20 million cells were plated on 75 mm^2^ flasks (non-ventilated cap) coated with poly-D-lysine and incubated at 37°C in 10% CO_2_. On DIV 3, non-astrocyte cells were removed by forceful shaking of closed flasks. Fibroblast elimination was performed by adding AraC to the media on DIV 5. On DIV 7, astrocytes were trypsinized (0.05% Trypsin-EDTA) and plated into 12-well (200,000 cells/well) or 6-well (400,000 cells/well) plates. On DIV 8, cultured rat astrocytes were transfected with shRNA plasmids using Lipofectamine LTX with Plus Reagent (Thermo Scientific) per the manufacturer’s protocol. Briefly, 1 µg (12-well) or 2 µg (6-well) total DNA was diluted in Opti-MEM containing Plus Reagent, mixed with Opti-MEM containing LTX (1:2 DNA to LTX) and incubated for 30 minutes at room temperature. The transfection solution was added to astrocyte cultures and incubated at 37C for 3 hours, then replaced with AGM. On DIV 10, astrocytes were trypsinized, resuspended in NGM, plated (20,000 cells per well) onto DIV 10 neurons, and co-cultured for 48 hours.

#### HEK293T

HEK293T cells used to produce lentivirus and adeno-associated virus were cultured in DMEM (GIBCO 11960) supplemented with 10% FBS, 100 U/mL Pen/Strep, 2 mM L-Glutamine, and 1mM sodium pyruvate. Cells were incubated at 37°C in 5% CO2 and passaged every 2-3 days.

### Plasmids

pLKO.1 Puro plasmids containing shRNA against mouse/rat *Ptprz1* were obtained from the RNAi Consortium (TRC) via Dharmacon; Clone ID: TRCN0000081069; shRNA sequence: CTCCTTAAACAGTGGCTCTAA. A scrambled shRNA sequence was generated by annealing the following oligonucleotides:

Fwd: CCGGGATAACCGTATTCACGCTATCCTCGAGGATAGCGTGAATACGGTTATCTTTTTG Rev: AATTCAAAAAGATAACCGTATTCACGCTATCCTCGAGGATAGCGTGAATACGGTTATC and cloned into the pLKO.1 Puro TRC cloning vector at AgeI and EcoRI restriction sites according to Addgene protocols and as described previously^38^. pLKO.1 shRNA plasmids expressing CAG-GFP in place of the Puromycin resistance gene were generated by restriction enzyme cloning at KpnI and SpeI sites as described previously^38^.

### Lentivirus production and transduction

Lentiviruses containing shRNA targeting vectors were produced by transfecting HEK293T cells with pLKO.1 shRNA-Puro, VSVG, and dR8.91 using X-tremeGENE (Roche). The following day, media was replaced with AGM and media containing lentivirus was collected on days 2 and 3 post-transfection. To test the knockdown efficiency of *Ptprz1* shRNA, DIV 7 rat primary cortical astrocytes were trypsinized and plated into 6-well plates (400,000 cells/well) in 2 mL of AGM. On DIV 8, 1 mL of AGM was removed from the astrocytes and replaced with 1 mL of media containing: 500 µL fresh AGM, 500 µL lentivirus-containing media, and 1 µg/mL polybrene. Cultures were treated with puromycin (1 µg/mL) from DIV 10-15 to eliminate non-transduced ells On DIV 15, protein was extracted using membrane solubilization buffer (MSB: 25 mM Tris pH 7.4, 150 mM NaCl, 1 mM CaCl2, 1 mM MgCl2, 0.5% NP-40, and protease inhibitors).

### Immunocytochemistry

Astrocyte-neuron co-cultures were fixed and stained as described previously^38^. Briefly, DIV 12 co-cultures were incubated with warm 4% PFA for 7 minutes, washed 3 times with PBS, and blocked in PBS containing 50% normal goat serum (NGS) and 0.4% Triton X-100 for 30 minutes at room temperature. Samples were washed once more in PBS and incubated overnight at 4°C with chicken anti-GFP (Aves GFP1020, 1:1000) diluted in antibody blocking buffer (ABB: pH 7.4, 150 mM NaCl, 50 mM Tris, 1% BSA, 100 mM L-lysine, 0.04% sodium azide) containing 10% NGS. The following day, samples were washed 3x with PBS, incubated with goat anti-Chicken 488 (Life Technologies, 1:500) diluted in ABB with 10% NGS for 2 hours at room temperature, and washed again 3x in PBS. Coverslips were mounted onto glass slides using Vectashield mounting media with DAPI (Vector Labs) and sealed with nail polish. Healthy astrocytes with strong expression GFP, a single nucleus, and minimal overlap with other GFP+ astrocytes, were imaged at 40x magnification in green and DAPI channels using Zeiss AxioImager M1. The individual acquiring the images was always blinded to the experimental condition. Sholl analysis was performed in FIJI as described previously^38^ and statistical analysis performed in RStudio using a linear mixed model with Tukey’s HSD^56^. At least 20 cells were imaged per condition per experiment, from three independent experiments. Cultures in which the peak complexity of the control condition was below 18 were excluded from analysis due to suboptimal cell health. Astrocytes containing multiple nuclei, weak GFP expression, or in areas of dense GFP labeled cells were not imaged.

### AAV Production and administration

To produce purified adeno-associated virus (AAV) pZac2.1-gfaABC1D-GFP-CAAX HEK293T cells were transfected with pAD-DELTA F6, serotype plasmid AAV PHP.eB, and pZac2.1-gfaABC1D-GFP-CAAX. Three days after transfection, cells were collected and lysed, and AAV-enriched fraction isolated from the supernatant by Optiprep density gradient and ultracentrifugation as described previously^57^. To label astrocytes with GFAP-CAAX, 1 µL of AAV containing pZac2.1-GfaABC1D-GFP-CAAX was injected unilaterally into the cortex of hypothermia-anesthetized P2 neonates using a Hamilton syringe.

### Tamoxifen administration

Tamoxifen administration was performed at P2 and P3 as described previously^38^. Briefly, tamoxifen powder was dissolved in corn oil at 10 mg/mL and further diluted in corn oil to 1.25 mg/mL (for P2 injection) and 2.5 mg/mL (for P3 injection). 40 µL of the respective tamoxifen solution was injected into the milk spot using an insulin syringe, for a dose of 0.05 mg at P2 and 0.1 mg at P3.

### Immunohistochemistry

#### Sample preparation

Brains were collected, frozen, sectioned, and stained as described previously^41^. Breifly, mice were anesthetized with 0.8 mg/kg tribromoethanol (avertin) and perfused with TBS/Heparin, followed by 4% PFA in TBS. Brains were post-fixed overnight in 4% PFA, rinsed 3x with TBS, and cryoprotected in 30% sucrose. Brains were frozen in embedding molds using a medium containing two parts 30% sucrose and one part O.C.T., and stored at -80 °C. Frozen brains were sectioned coronally to 25, 40, or 100um thickness on a CryoStar NX50 Cryostat (Thermo Fisher Scientific) and stored in a 1:1 mixture of glycerol and 1xTBS at -25 °C until use. For immunostaining, sections were washed in TBST (0.2% Triton in TBS), blocked in blocking solution (10% goat serum in TBST), and incubated in primary antibody solution (primary antibody diluted in blocking solution) for 2-3 nights at 4 °C while shaking at 100 rpm (see specific sections below for antibody concentrations). Following primary antibody incubation, sections were washed in TBST, incubated in secondary antibody solution (secondary antibody diluted 1:200 in blocking solution) for 2-3 hours at room temperature, and washed again in TBST. DAPI was added to the secondary antibody solution for the final 10 minutes of incubation at a 1:50,000 concentration. Sections were then mounted onto glass slides with homemade mounting medium (20 mM Tris pH 8.0, 90% Glycerol, 0.5% N-propyl gallate) and sealed with nail polish. For primary antibodies produced in mouse, isotype specific secondary antibodies were used (e.g., goat anti-mouse IgG1) to prevent excessive background staining.

#### Cell counting

40 µm-thick sections containing visual cortex were labeled with one of two antibody combinations: 1) Rb Sox9 (Millipore AB5535, 1:1000), Ms IgG2a Olig2 (Millipore MABN50, 1:400), and DAPI, or 2) Ms IgG1 NeuN (Millipore MAB377, 1:1000), and DAPI. Corresponding secondary antibodies produced in goat (Life Technologies) were used at 1:200. Tile scan images were acquired from P21 *Ptprz1* control (WT and cHet) and cKO mice using an Olympus FV3000RS inverted confocal microscope with a resonant scanner and 20x objective. For each brain section, an ROI of size 447.47 μm x 930.98 μm spanning L1 through L6 of the visual cortex was selected for analysis of cell number. All image processing and analysis was completed using novel, semi-automated CellProfiler pipelines (available at https://github.com/BaldwinLabUNC/Astrocyte_morphology). Images were denoised using the GaussianFilter module, then nuclei signal was enhanced and background signal was suppressed using the EnhanceOrSuppressFeatures module. DAPI+ nuclei were identified with the IdentifyPrimaryObjects module and new images with the identified nuclei were saved. To identify Sox9+ and Olig2+ nuclei, the EnhanceEdges module was used to improve the identification of nuclei and help distinguish nuclei from debris, then objects were segmented using the IdentifyPrimaryObjects module. Mean fractional intensities (MeanFrac) of the objects were measured using the MeasureObjectIntensityDistribution module, and a single maximum MeanFrac value was used for each animal in the FilterObjects module as a threshold to filter out non-nuclear objects. New images with the identified nuclei were saved, then co-localized Sox9+ and Olig2+ nuclei were identified using the RelateObjects module and saved as new images. To identify NeuN+, objects were segmented using the IdentifyPrimaryObjects module, then objects were filtered to exclude non-nuclear objects using the FilterObjects module based on eccentricity values measured in the MeasureObjectSizeShape module, and new images with the identified nuclei were saved. In all cases, the ExportToSpreadsheet module was used to count the number of objects in the saved images containing identified nuclei, and these counts were used as the Cells/Image data points. For each staining condition, 3 sections per brain from 6 sex-matched littermate pairs were collected and analyzed. Differences in cell number between genotype were analyzed using paired two-tailed t test. The experimenters were blinded to subject group during image acquisition and Imaris analysis. All mice that appeared healthy at the time of collection were included in this study. No data were excluded.

#### Astrocyte 3D morphology analysis

Individual astrocyte territory volume and surface area was assessed in 100μm-thick floating sections of mouse primary visual cortex collected at postnatal day 21 (P21). Tissue from Control and Ptprz1 cKO mice with intracortical AAV injection of GFP-CAAX was collected, processed, and stained as described above using chicken anti-GFP primary antibody (Aves GFP1020, 1:1000), goat anti-chicken Alexa Fluor 488 secondary antibody (1:200), and DAPI (1:50,000). High magnification images containing whole astrocytes (50-60μm z-stack) were acquired on an Olympus FV3000 microscope with a 40x objective and 2x optical zoom. Inclusion criteria for analysis required the entirety of the astrocyte to be contained within a single brain section, specifically visual cortex layer 5. Astrocytes outside of this brain region and/or incomplete astrocytes were excluded from this study. Imaris (Bitplane) software was used as described previously to analyze astrocyte territory volume^41^. Briefly, minimal post-processing (Median Filter 3×3×3, Background Subtraction sigma=40.00, Normalize Layers) was performed through batch processing in Imaris to aid whole-cell surface reconstruction with the surface creation tool. Spots close to surfaces were then generated and used by the custom Convex Hull Xtension^41^ to build a convex hull around the whole astrocyte territory. A small number of astrocytes were excluded from the final analysis due to exceptionally dim fluorescent signal or poor subject tissue and image quality. Files containing detailed surface (astrocyte) and convex (territory) metrics were exported from Imaris for each individual astrocyte. A custom RStudio script (available at: https://github.com/BaldwinLabUNC/Astrocyte_morphology) was developed to extract appropriate descriptive metrics (Area, Oblate Ellipticity, Prolate Ellipticity, Sphericity, and Volume) from exported CSV files, combine data into a single CSV file for additional statistical analysis in GraphPad Prism 10, and run preliminary statistical analyses on animal averages: Shapiro-Wilk normality test, F-test, and subsequently appropriate unpaired two-sample t-tests (Student’s t-test and Mann-Whitney U test). Normality and F-test results were used to determine appropriate t-tests run in GraphPad Prism 10 for plotting and P-value reporting. Statistics using individual cell values as opposed to animal average values were conducted in GraphPad Prism 10 using the same above-mentioned tests for normality and variance prior to conducting appropriate unpaired two-sample t-tests. The experimenter was blinded to subject group during image acquisition and analysis in Imaris. The number of animals and cells/animal analyzed is indicated in the corresponding figure legend for this experiment.

#### Neuropil infiltration volume analysis

Astrocyte infiltration into the surrounding neuropil was analyzed as described previously^38^ in 40μm-thick floating sections of P21 mouse primary visual cortex. High magnification Z-stack images were acquired on an Olympus FV3000 microscope with a 60x objective at 2x optical zoom. Astrocytes were sparsely labeled with GFP-CAAX and *Ptprz1* was knocked down in applicable subjects as described above. Inclusion criteria required astrocytes be located in layer 5 of the visual cortex, demonstrate sufficiently bright fluorescent labeling, encompass the entire astrocyte arbor in the X/Y plane, and include at least 10 μm of the astrocyte arbor above and below the soma in the Z-stack. Astrocytes failing to meet these criteria were not imaged or otherwise excluded from the final analysis. For each astrocyte, three ROIs (12.65 μm x 12.65 μm x 10um) containing only neuropil (excluding astrocyte soma, large branches, and endfeet) were selected and reconstructed in Imaris using the surface tool. Surface volume within individual ROIs was recorded and averaged for each cell (3 x ROIs) and animal (3 x ROIs across 4-5 cells per animal). Additional astrocytes were excluded from the final analysis for being a duplicate (one cell) or twin astrocyte (one cell), decreasing the total cells from 5 to 4 for two subjects. The experimenters were blinded to subject group during image acquisition and Imaris analysis. Data from biological replicates were analyzed using a nested t-test. Animal averages were analyzed using an unpaired two-sample t-test (Student’s t-test). All statistical analyses were conducted in GraphPad Prism 10 and the number of animals and cells/animal analyzed is indicated in the corresponding figure legends for this experiment.

#### Astrocyte 2D morphology analysis

High-magnification Z-stack images as acquired for NIV analysis were subsequently used to conduct 2D morphology analysis as previously described^42^. Images were flattened in FIJI/ImageJ software through maximum intensity Z-projections. Individual brightness levels were adjusted for each cell to enable selection of the complete astrocyte territory with the magic wand selection tool (8-connected mode). Astrocyte morphology measurements included Feret’s max and min diameter, aspect ratio, territory area, circularity, and roundness as described in the figure and figure legend for this experiment. The experimenter was blinded to subject group during FIJI/ImageJ analysis. Animal averages were analyzed using appropriate tests (unpaired two-sample t-test or Mann-Whitney U test). All statistical analyses were conducted in GraphPad Prism 10 and the number of animals and cells/animal analyzed is indicated in the corresponding figure legends for this experiment.

#### Synapse imaging and analysis

Synaptic staining was performed in 25 μm-thick coronal sections containing the visual cortex from P21 *Ptprz1* control (WT and cHet) and cKO mice. Three different antibody combinations consisting of a presynaptic and postsynaptic target were used to label three different types of synapses: 1) excitatory intracortical: VGlut1 (Synaptic Systems 135 304, 1:1000) and PSD95 (Thermo Fisher 51-6900, 1:300), 2) excitatory thalamocortical: VGlut2 (Synaptic Systems 135 404, 1:2000) and PSD95, and 3) inhibitory: VGAT (Synaptic Systems 131 004, 1:1000) and Gephyrin (Synaptic Systems 147 021, 1:200). Corresponding Alexa Fluor-conjugated secondary antibodies produced in goat (Life Technologies) were used at 1:200. For VGlut1/PSD95 imaging, high magnification 63x objective z-stack images containing 15 optical sections spaced 0.34 μm apart were obtained using the Leica SP8X Falcon inverted confocal microscope. For VGlut2/PSD95 and VGAT/gephyrin, z-stack images containing 15 optical sections spaced 0.34 μm apart were acquired with a 60x objective and 1.64x optical zoom using an Olympus FV3000. Co-localized presynaptic and postsynaptic puncta were quantified using Synbot^49^ with the following parameters: 2-channel, noise reduction, manual thresholding, minimum pixel = 3, pixel overlap. For each staining condition, 6 sex-matched littermate pairs were collected and analyzed. For each animal, 3 z-stack images (15 slices) were acquired and each image converted in 5 separate maximum projection images (MPI) of 3 slices each for a total of 15 MPIs per animal. Animal averages were analyzed via paired two-tailed student’s t test using Graphpad Prism 10. The individual acquiring the images and performing the analysis was always blinded to the experimental condition. No data were exclueded from the analysis.

### Protein extraction and western blotting

For analysis of cortical lysates via western blot, *Ptprz1* WT, cHet, and cKO mice were anesthetized with 0.8 mg/kg tribromoethanol (avertin) and perfused with TBS/Heparin to minimize IgG contamination from blood. Immediately following perfusion, brain cortices were rapidly dissected, and flash frozen in liquid nitrogen. Brain lysis was performed as described previously^38^. Briefly, one half of the cortex was homogenized in 1 mL of Lysis Buffer R (150 mM NaCl, 50 mM Tris, 1mM EDTA, pH 7.5, protease inhibitor mix, 1 mM Na_3_VO_4_, 20 mM NaF, and 10 mM beta-glycerophosphate) using a ceramic pestle and glass tube. Homogenate was collected and combined with an equal volume of modified RIPA buffer lacking SDS (M-RIPA: 50 mM Tris, 150 mM NaCl, 1 mM EDTA, 2% NP40, 2% deoxycholate, pH 7.5 containing protease inhibitors (Roche) 1 mM Na3VO4, 20 mM NaF, and 10 mM beta-glycerophosphate). Lysis proceeded for 20 minutes with rotation at 4°C (12-15 rpm), then the lysate was centrifuged at max speed at 4°C for 10 minutes. Supernatant was collected and the Pierce BSA Protein Assay Kit (Thermo Fisher) was used to determine protein concentration. Samples were stored at -80°C until use.

Chondroitinase ABC lysate digest was performed to remove glycosaminoglycan side chains from PTPRZ1. 10 U Chondroitinase ABC (Sigma-Aldrich) was reconstituted in reconstitution buffer (1% BSA, 50 mM Tris pH 8.0) and stored at -80°C until use. For the digest, a 2 U/mL chondroitinase ABC working solution was prepared using dilution buffer (2% BSA, 60 mM sodium acetate, 50 mM Tris pH 8.0) and combined with an equal volume of lysate containing 100 µg. Samples were incubated at 37°C for 90 minutes then mixed with 4x Laemmli Sample Buffer (Bio-Rad) containing 5% β-ME and incubated for 45 minutes at 45°C for denaturation. 20 μg of protein was loaded into 4%-15% gradient pre-cast gels (Bio-Rad) and run at 50V for 5 minutes followed by 150 V for 1 hour. Proteins were transferred to PVDF membrane (Millipore) at 100 V for 2 hours. Immediately following the transfer step, Method 1 provided by the LI COR BIOTECH LLC Revert^TM^ 700 Total Protein Stain Kit for Western Blot Normalization was performed to obtain a total protein control stain. Briefly, the PVDF membrane was fully dried, rehydrated in 100% methanol, incubated in Revert 700 Total Protein Stain solution, and immediately imaged on a LI-COR Odyssey imaging system. Following total protein detection, the membrane was blocked in Intercept Blocking Buffer (LI-COR) and incubated in primary antibody overnight at 4°C (PTPRZ Rabbit anti-Human, Mouse, Rat Polyclonal Antibody, 1:1000, Fisher Scientific). The next day, the membrane was washed with TBS-Tween-20, incubated in LI-COR secondary antibody solution (Anti-Rabbit IgG Goat Secondary Antibody, IRDye® 800CW, 1:5000, Avantor) for 2 hours with agitation at ambient temperature, washed twice with TBS-Tween-20 and once with TBS, then immediately imaged on a LI-COR Odyssey imaging system. Protein expression was quantified using Image Studio Lite software.

### Quantification and Statistical Analysis

All statistical analyses were performed in GraphPad Prism 10, with the exception of the Sholl analysis experiments where statistical analysis was performed in RStudio as described previously^56^. For each experiment, the number of subjects and specific statistical tests are included in the figure legend and data are represented as mean ± standard error of the mean with exact P-values shown. Sample sizes were determined based on previous experience for each experiment and no statistical methods were used to predetermine sample size. Specific details for inclusion, exclusion, and randomization are included in specific method subsections.

## Data and code availability

All custom code available at https://github.com/BaldwinLabUNC/Astrocyte_morphology. Data available upon request.

## Author Contributions

A.R.E: conceptualization, methodology, investigation, formal analysis, visualization, writing – review and editing. H.E.S: methodology, investigation, formal analysis, software, visualization, writing – review and editing. K.T.B: conceptualization, methodology, investigation, formal analysis, supervision, funding acquisition, visualization, writing – original draft.

## Acknowledgments

We thank Dr. Cagla Eroglu for providing the *Ptprz1* floxed mouse line, and the Duke Transgenic Mouse Facility for generating the *Ptprz1* floxed mouse line. Microscopy was performed at the UNC Neuroscience Microscopy Core (RRID:SCR_019060), supported, in part, by funding from the NIH-NICHD Intellectual and Developmental Disabilities Research Center Support Grant P50 HD103573. The Baldwin Lab is supported by the NIH, DP2 NS136873 to K.T.B. and T32NS007431 to H.E.S.

## Data Availability Statement

Data available upon request. All custom code available at https://github.com/BaldwinLabUNC/Astrocyte_morphology.

## Ethics Statement

All experimental protocols were performed in accordance with NIH guidelines and received approval from the Animal Care and Use Committee of UNC Chapel Hill.

---

**Supplementary Figure 1:**
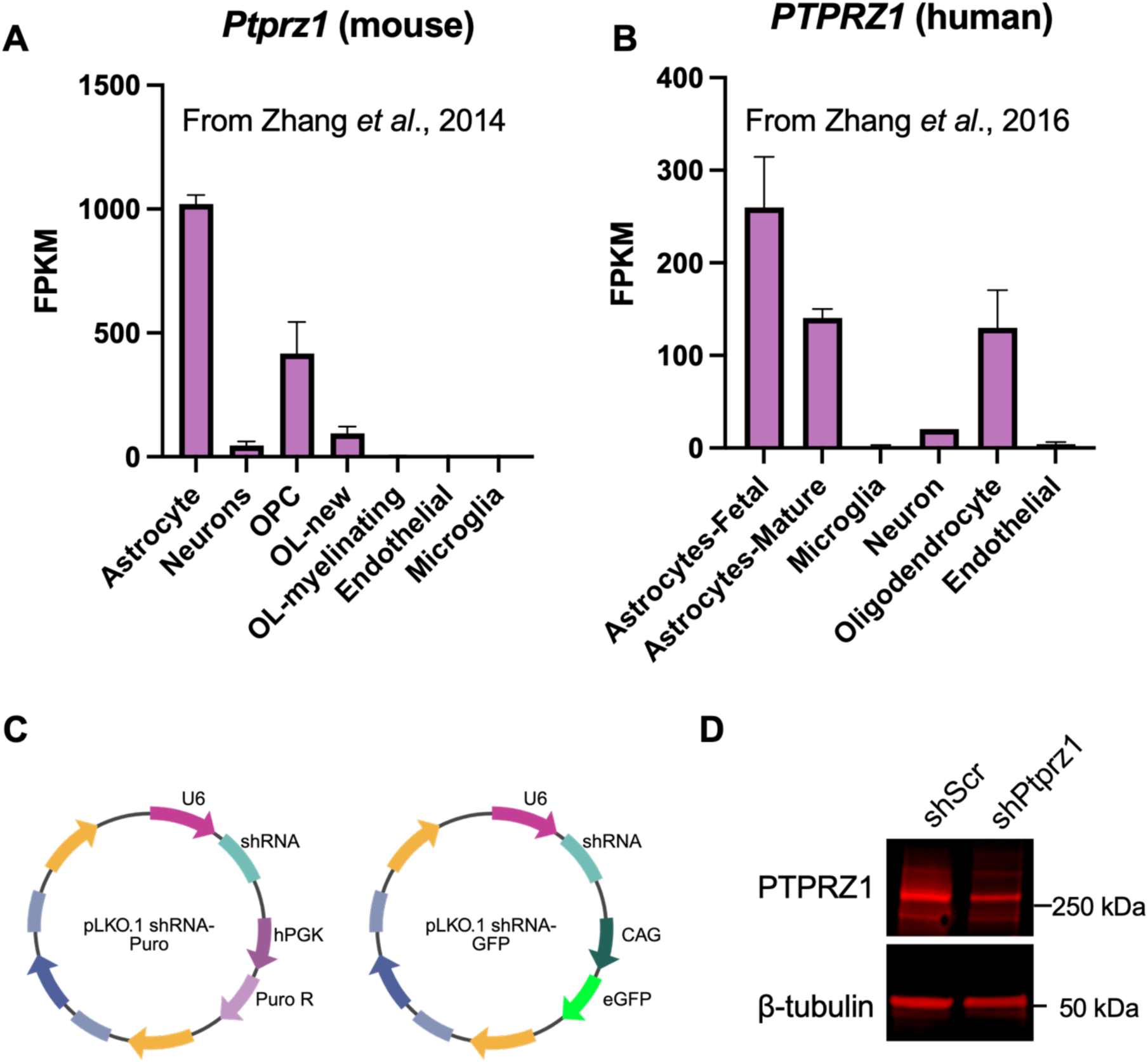
Ptprz1 expression and shRNA validation. **A)** *Ptprz1* gene expression levels per cell type in mouse from Zhang et al., 2014^5^. **B)** PTPRZ1 gene expression levels per cell type in human from Zhang et al., 2016^6^. **C)** Plasmid maps for pLKO.1 vectors used in this study. Plasmids expressing shRNA and puromycin resistance (PuroR) were packaged into lentivirus and transduced into astrocytes to validate shRNA knockdown efficiency (pLKO.1 shRNA-Puro). For morphology analysis, the hPGK promoter and Puro R were replaced with a CAG promoter driving expression of eGFP (pLKO.1 shRNA-GFP). Maps created with BioRender.com. **D)** Western blot of primary rat astrocytes transduced with lentivirus expressing pLKO.1 shScr-Puro or pLKO.1 shPtprz1-Puro and treated with puromycin to eliminate non-transduced astrocytes. PTPRZ1 labeling demonstrates effective knockdown with shPTPRZ1. β-tubulin is used as a loading control.

**Supplementary Figure 2:**
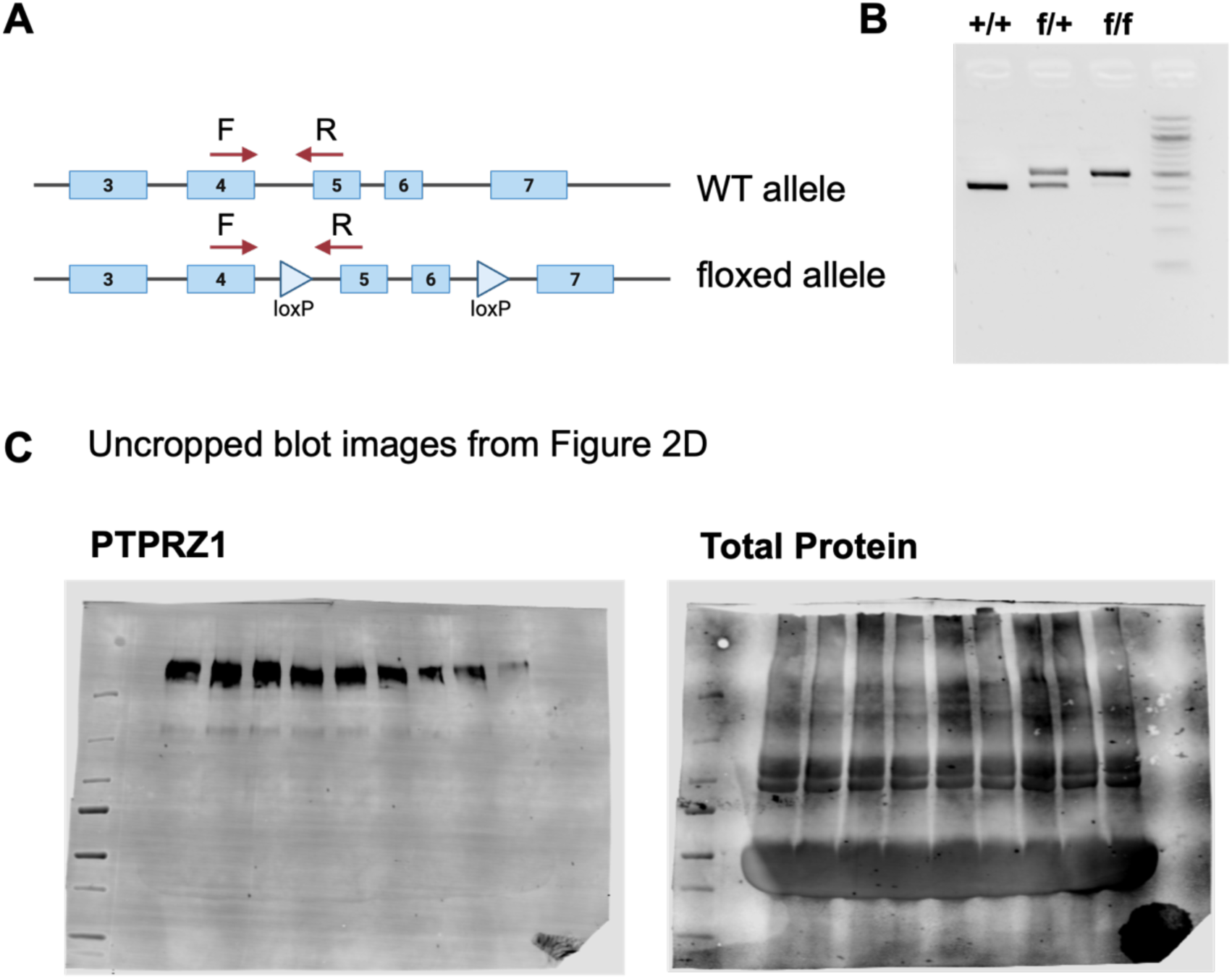
cKO genotyping strategy. **A)** Genotyping strategy for detecting wild-type (WT) and floxed *Ptprz1* alleles from mouse genomic DNA. The same forward and reverse primers detect both alleles, with the floxed allele appearing 89 base pairs higher due to the addition of the *loxP* site. **B)** Example of PCR products obtained from WT (+/+), f/+, and f/f mice. **C)** Uncropped western blot images from Figure 2D.

**Supplementary Figure 3:**
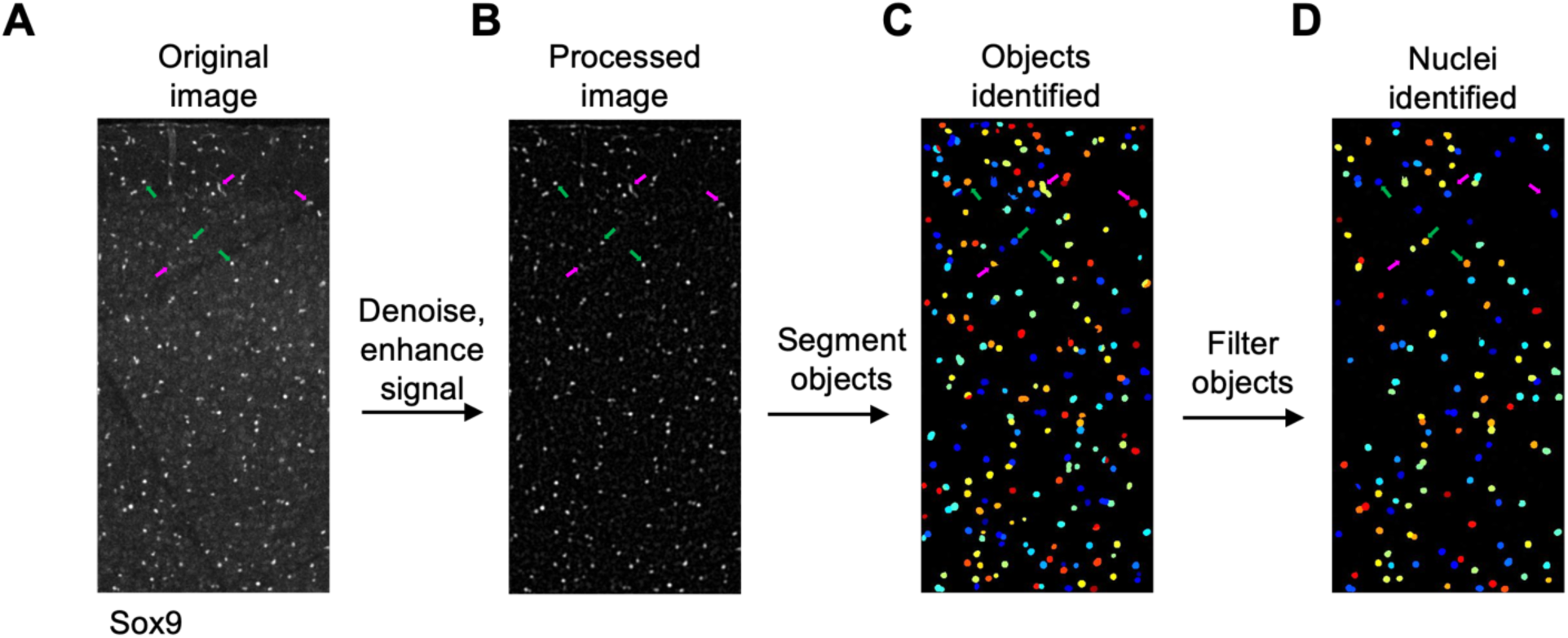
Cell counting workflow. **A)** Representative input image for the cell count pipeline. Cropped grayscale image of visual cortex with Sox9-labeled nuclei. Green arrows denote representative Sox9+ nuclei that will eventually be included in the cell count. Magenta arrows denote representative debris that will eventually be excluded from the cell count. **B)** Processed input image, after denoising, foreground signal enhancement, and nuclei-specific signal enhancement. Signal separates more clearly from the background, and nuclei appear more distinct from debris, compared to the original image. **C)** Identified objects segmented from the processed image. Both nuclei and debris are identified as objects. **D)** Objects representing debris are filtered out. Objects representing Sox9+ nuclei remain and are included in the quantification of Sox9+ cells in the image.

**Supplementary Figure 4:**
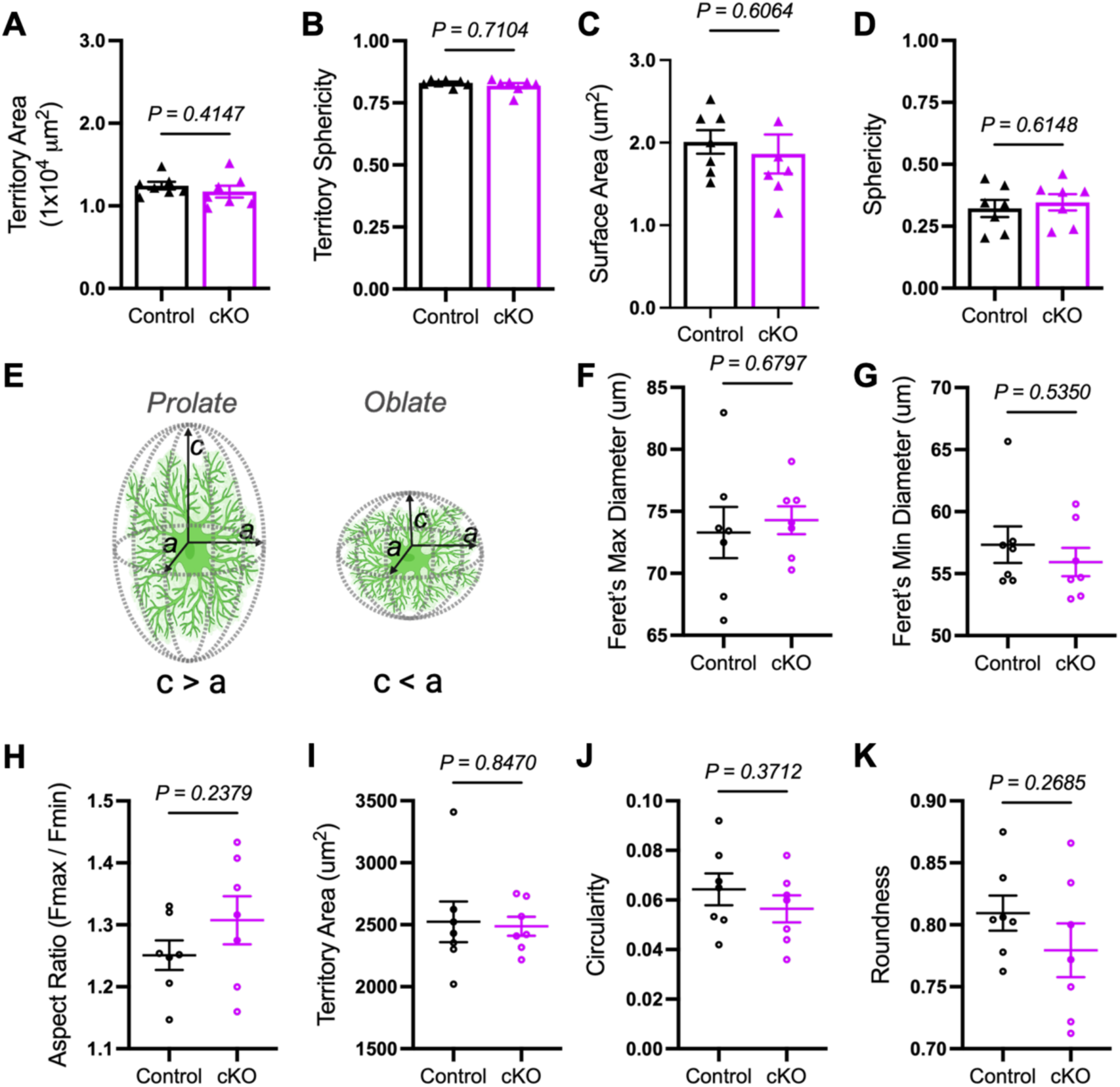
Additional morphological analyses. **(A-D)** Additional 3D morphology analyses of: **(A)** territory area, **(B)** territory sphericity, **(C)** astrocyte surface area, and **(D)** astrocyte surface sphericity in P21 V1 L5 astrocytes. Data presented as animal averages (left; individual mice represented as triangles; n=7 mice per group, 4-8 cells per mouse) and individual astrocyte statistics (right; individual astrocyte represented by open circles). Bars are mean +/- SEM. Unpaired t-test (A, C-D) or Mann-Whitney U test (B). **E)** Schematic describing prolate vs oblate ellipticity as a difference in whether the ellipse is rotated about a major (longest) axis or minor (shortest) axis. **(F-K)** 2D morphology analyses of: **(F)** Feret’s max diameter, **(G)** Feret’s min diameter, **(H)** aspect ratio (Feret Max/Feret Min), **(I)** territory area, **(J)** circularity, and **(K)** roundness. Dots represent animal averages (n=7 mice per group, 4-6 cells per mouse). Bars are mean +/- SEM. Unpaired t-test (F, H-K) or Mann-Whitney U test (G).

**Supplementary Figure 5:**
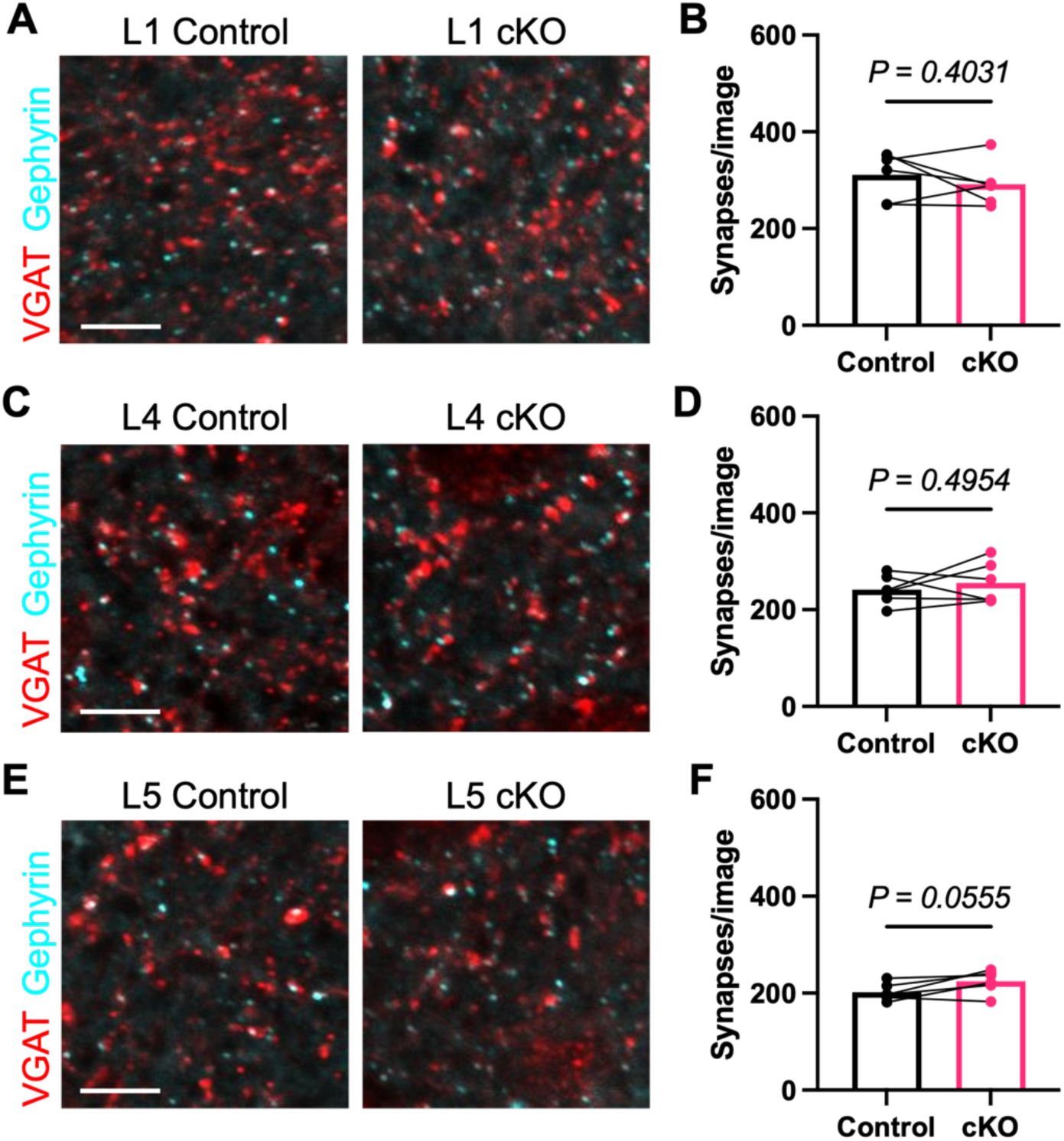
Inhibitory synapse number is unchanged at P21. **A)** Representative 20 µm x 20 µm regions of interest (ROIs) labeled with inhibitory synapse markers in L1 visual cortex at P21. Presynaptic VGAT (red) and postsynaptic marker gephyrin (cyan). Scale bar 5 µm. **B)** Quantification of number of co-localized VGAT/gephyrin puncta per image (129.35 µm x 129.35 µm) from L1 visual cortex. **C)** Representative ROIs labeled with VGAT and gephyrin in L4 and **D)** quantification of co-localized puncta. **E)** Representative ROIs labeled with VGAT and gephyrin in L5 and **F)** quantification of co-localized puncta. For B, D, and F: n = 6 sex-matched littermate pairs of control and cKO mice. Lines connect sex-matched control-cKO littermates. Dots represent per animal averages of 15 images. Paired two-tailed student’s t-test.

